# A *Salmonella enterica* serovar Typhimurium Genome-wide CRISPRi Screen Reveals a Role for Type 1 Fimbriae in Evasion of Antibody-Mediated Agglutination

**DOI:** 10.1101/2024.12.16.628777

**Authors:** Samantha K. Lindberg, Graham G. Willsey, Nicholas J. Mantis

## Abstract

The O5-specific monoclonal IgA antibody, Sal4, mediates the conversion of *Salmonella enterica* serovar Typhimurium (STm) from virulent, free-swimming cells to non-motile, multicellular aggregates in solution (“flocs”) as well as biofilm-like associations at the air-liquid interface (ALI). We hypothesize that the rapid transition from an invasive to a non-invasive state reflects an adaptation of STm to Sal4 IgA exposure. In this report, we performed a genome-wide CRISPRi screen to identify STm genes that influence multicellular aggregate formation in response to Sal4 IgA treatment. From a customized library of >36,000 spacers, ∼1% (373) were enriched at ALI after two consecutive rounds of Sal4 IgA treatment. The enriched spacers mapped to a diversity of targets, including genes involved in O-antigen modification, cyclic-di-GMP metabolism, outer membrane biosynthesis/signaling, and invasion/virulence, with the most frequently targeted gene being *fimW*, which encodes a negative regulator of Type 1 Fimbriae (T1F) expression. Generation of a STm Δ*fimW* strain confirmed that the loss of FimW results in a hyperfimbriated phenotype and evasion of Sal4 IgA-mediated agglutination in solution. Closer examination of the *fimW* mutant revealed its propensity to form biofilms at the ALI specifically in response to Sal4 exposure, suggesting that T1F “primes” STm to transition from a planktonic to a sessile state possibly by facilitating bacterial attachment to abiotic surfaces. These findings shed light on the mechanism by which protective secretory IgA antibodies influence STm virulence in the intestinal environment.

## INTRODUCTION

The intestinal epithelium is a point of entry for a multitude of enteric pathogens, which are collectively responsible for significant morbidity and mortality worldwide, especially in children under the age of five (1, 2). One pathogen of increasing concern is *Salmonella enterica* serovar Typhimurium (STm), a motile, facultative anerobic, Gram-negative bacterium typically associated with self-limiting gastroenteritis. However, the past fifty years has seen a rise in multidrug-resistant and host-adapted isolates of STm that are capable of causing invasive non-typhoidal *Salmonella* infections (3–8). Invasion of intestinal tissues by STm is a complex, multistep process involving flagellar-based motility and a variety of adhesins (e.g., fimbriae) to secure contact with the apical surfaces of intestinal epithelial cells (9). STm then employs a specialized Salmonella pathogenicity island-1 (SPI-1) encoded type-three secretion system (T3SS-1) along with an array of effector proteins involved in to gain entry into the host cells (9). After breaching the epithelial barrier, STm can disseminate systematically and infect multiple different organ types (10–12).

Immunoglobulin A (IgA) antibodies are actively transported into the intestinal secretions where they are capable of intercepting STm the intestinal lumen and preventing bacterial attachment to epithelial cells, in a process known as immune exclusion (13, 14). Immune exclusion involves antibody-mediated bacterial clumping, which occurs through two distinct mechanisms depending on local cell density (15, 16). So-called “enchained growth” occurs at low cell density (<10^7^ CFU/g), where IgA cross-links actively dividing cells and prevents their separation (17). At high cell density, neighboring cells are cross-linked via the formation of antibody-mediated intercellular bridges. This type of aggregation, referred to as “classical agglutination,” occurs only at cell densities where cell-cell contacts are frequent (16). Once initiated, classical agglutination of STm results in the formation of macroscopic bacterial mats that share many hallmarks associated with bacterial biofilms, including ECM production and loss of motility (18–22).

The mouse monoclonal IgA Sal4 promotes agglutination of STm, and this directly correlates with the ability of the antibody to limit invasion of epithelial cells *in vitro* and *in vivo* (23–28). Sal4 IgA targets the immunodominant O5 antigen (O-Ag) of lipopolysaccharide molecules expressed on the surface of STm (23, 29). The presence of an acetylated abequose moiety attached to the trisaccharide O-Ag backbone (O12) confers the O5-antigen phenotype (30). The acetyltransferase encoded by the *oafA* gene is required for O5^+^ antigen expression and thus Sal4 recognition (31). Michetti and colleagues observed that Sal4 IgA prevents the ability of wild type (WT) STm to invade confluent epithelial cell monolayers, but had no effect on a STm *oafA* mutant (24). Moreover, protection occurred at concentrations of Sal4 sufficient to induce visible bacterial agglutination and immune complex formation. In subsequent studies, it has been demonstrated that Sal4 IgA treatment of STm is accompanied by rapid motility inhibition, concurrent exopolysaccharide (EPS) production and the eventual formation of multicellular macroscopic aggregates (“flocs”) and biofilm-like mats on abiotic surfaces (e.g., borosilicate glass) (19, 20, 22, 25). In a mouse model, oral administration of Sal4 IgA entraps STm within the intestinal lumen and reduces bacterial invasion into gut-associated lymphoid tissues by several orders of magnitude, demonstrating the essential role Sal4-mediated agglutination plays in immunity against STm during infection (22).

We recently developed the so-called snow globe assay (SGA) to visualize and quantify Sal4-mediated classical agglutination of STm (32). This method utilizes high (∼10^8^ CFU/mL) cell density cultures, either of individual strains or mixed populations, to directly examine classical agglutination of STm caused by the combination of antibody cross-linking and cell-cell collisions. We investigated the role of flagellar-based motility in promoting antibody-mediated agglutination and found that bacterial cell collisions drive this process rather than motility per se. The role of several cyclic-di-GMP metabolizing enzymes in contributing to Sal4-mediated agglutination was investigated as well, with no observable phenotypic differences relative to a WT control in the handful of mutants tested (32).

Based on these findings, the goal of the current study was to identify genetic components in STm that modulate the extent of Sal4-mediated agglutination of STm. We used a previously generated CRISPR interference (CRISPRi) library in STm strain 14028s to screen for genes that influence agglutination in the presence of Sal4 IgA. The screen revealed that repeated passages of Sal4 IgA treatment enriched for spacers that mapped to the *fimW* locus. The gene product of *fimW* is a negative regulator of T1F expression, and this function was confirmed with a fully quantitative mannose-sensitive yeast agglutination assay. Further characterization of a STm *fimW* mutant strain in the presence of Sal4 demonstrated that enhanced T1F expression facilitates the transition to a biofilm-like state by promoting adherence to glass and polystyrene at the ALI following antibody exposure. Understanding how a host immune factor like IgA can drive pathogen adaptation has implications for advancing IgA mAbs as mucosal therapeutics to prevent bacterial persistence in the GI tract.

## RESULTS

### A STm genome-wide CRISPRi screen identifies Sal4 IgA “escape” mutants

Based on our previous investigations of the role of motility and cyclic-di-GMP metabolism in driving Sal4 IgA-mediated agglutination, we set out to identify additional STm genes whose expression influences this process (32). To accomplish this, we performed a CRISPR interference-based screen (CRISPRi) of STm 14028s using a previously generated library of 36,651 plasmid-based single guide RNAs, or spacers, which target promoters and genes across the chromosome and virulence plasmid (J. Wade, unpublished) (33). Multiple (up to six) spacers were designed for each target to account for potential off-target and/or ineffective DNA binding. The STm strain containing the library (SL061; Table S1) also features a Δ*cas3::thyA* mutation that enhances expression of the downstream *cas* operon and inhibits endonuclease activity, thus enabling the endogenous Type I-E CRISPR-Cas system to repress self-gene expression in a customizable, sequence-specific manner (34). The 32-nucleotide-long library spacers were designed relatively uniformly across the STm genome, theoretically avoiding bias towards larger gene targets that can be encountered with transposon mutagenesis (35).

We leveraged the SGA to screen for spacers that were over- and under-represented at the air-liquid interface (ALI) after repeated Sal4 IgA exposure. We reasoned that hypo-agglutinating cells would be enriched at the ALI, while hyper-agglutinators would be de-enriched at the ALI. As a proof of concept that we could enrich for targets of interest, we spiked a culture of strain SL174, a kanamycin-resistant derivative of STm 14028s constitutively expressing b-galactosidase (O5^+^; *lacZ^+^*) with a STm Δ*oafA* (O5*^-^*; *lacZ^-^*) strain at a ratio of 6000:1 and subjected the mixture to four rounds of Sal4 IgA (15 µg/mL) treatment (Table S1). Following 5 h of treatment, aliquots were taken from the ALI and transferred into fresh medium to grow overnight and continue the enrichment the following day. In parallel, aliquots were spotted onto LB agar containing X-gal (5-bromo-4-chloro-3-indolyl-β-D-galactopyranoside) after each treatment period to monitor the ratios of blue (WT-*lacZ*) and white (STm Δ*oafA*) colonies. Four consecutive rounds of Sal4 IgA treatment enriched for the *oafA* mutant, which ultimately comprised >80% of the culture (**Figure 1**). In the absence of Sal4 IgA treatment, the *oafA* mutant was undetectable even after four passages. These results confirm previous observations that serial treatment of STm with Sal4 IgA enriches for Sal4 escape mutants (23, 29).

**Figure 1:**
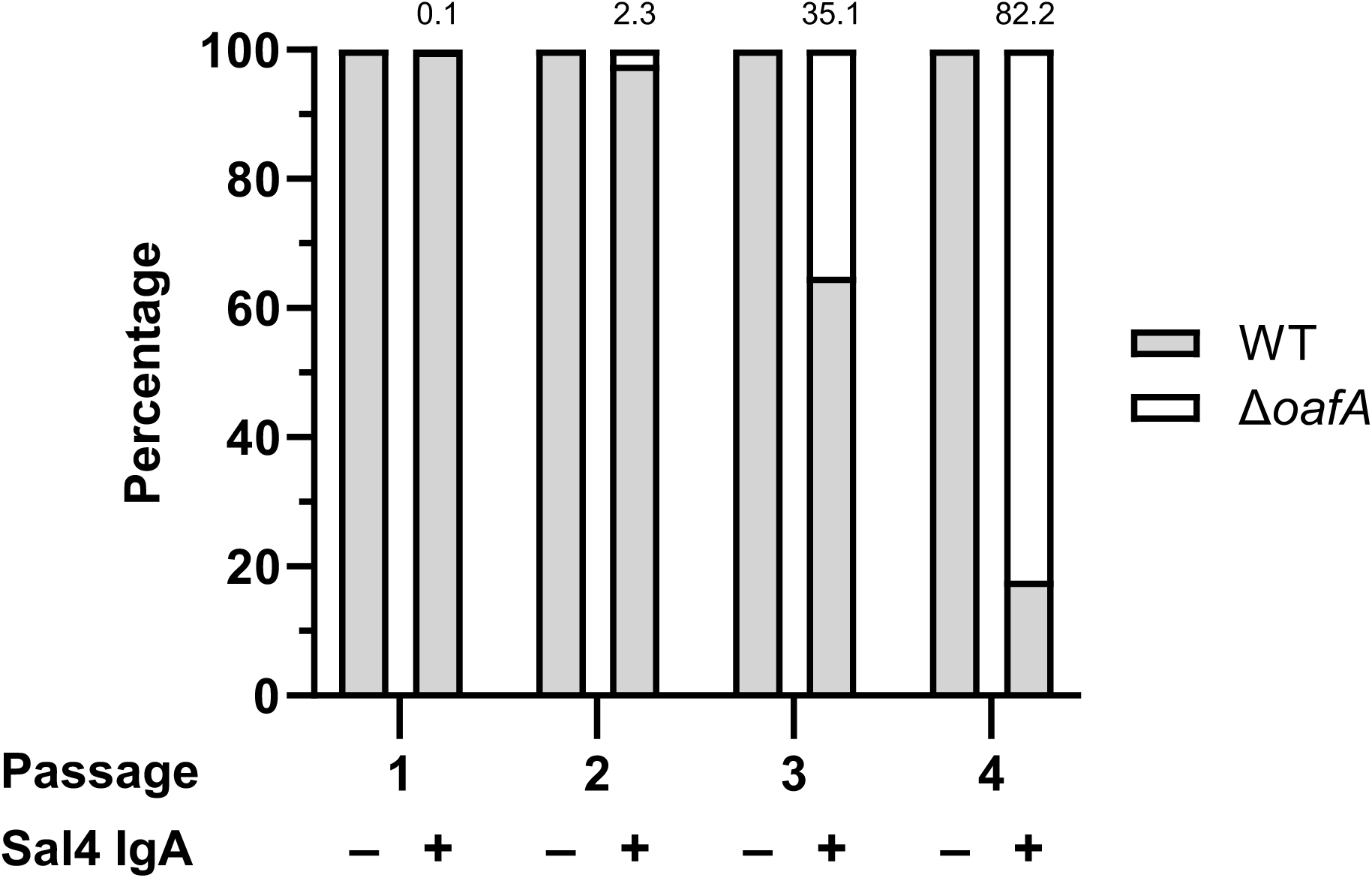
Repeated Sal4 IgA treatment enriches for O5^-^ STm in a predominantly O5^+^ population. Mid-log phase cultures of WT (SL174; *lacZ*^+^) and Δ*oafA* (SL180; *lacZ*^-^) STm were washed in PBS, combined at a ratio of approximately 6000:1, and then left untreated or treated with 15 μg/mL of Sal4 IgA. At 5 h post-treatment, 100 µL of the culture from the top of the supernatant was collected and plated on LB agar containing X-gal to determine the relative composition of each strain. This portion of the culture was passaged, and the assay procedure was repeated the following day for a total of four rounds of treatment. Data represents the percentage of each strain (with the value for Δ*oafA* listed above the bar for the Sal4-treated groups) averaged from two biological replicates each with two technical replicates.

To screen for additional escape mutants, we subjected the STm CRISPRi library to two consecutive passages in the absence (control) or presence of Sal4 IgA (15 µg/mL) treatment. Following the second round, an aliquot (200 µL) was taken from the ALI, cultured overnight, and the spacer-containing plasmids were purified. The spacer elements were PCR amplified in bulk and the resulting amplicons were submitted for Next-Generation Sequencing (NGS) with the Illumina NextSeq platform. A Python script was used to determine the frequency of each spacer in the control and Sal4-treated cultures from the raw .fastq files (Dataset S2). We then developed and implemented a custom R script to establish a minimum frequency of 100 reads per spacer, calculate the fold change in spacer frequency between treatment conditions (untreated versus Sal4 IgA-treated), and then map the spacers to their corresponding target gene with RefSeq annotations (Dataset S3). A spacer was classified as enriched if its frequency in the Sal4-treated cultures was >2-fold over that in the untreated cultures (see **Figure S1** for complete analysis workflow).

The screen yielded a total of 373 spacers that were enriched at the ALI after two rounds of Sal4 IgA treatment (**Dataset S4**). These spacers mapped to 307 genes and 47 promoters associated with range of known or predicted gene products, including cyclic-di-GMP metabolism (*yfeA*; *yfiN*), outer membrane composition (*yhfL*; *ompC*), outer membrane response signaling (*ompR*; *envZ*), chemotaxis (*cheR*), and invasion/virulence (*invA*). Sixteen genes were each targeted by 2 enriched spacers and two genes were targeted by 3 unique spacers, one of which was *oafA,* thereby validating the CRISPRi screen for the ability to identify escape mutants. The other gene targeted by three spacers was *fimW*, which encodes a negative regulator of the Type 1 Fimbriae (T1F) operon (36). Interestingly, three additional unique spacers targeted the upstream region of *fimW* (formerly annotated as *stm14_0645*), possibly overlapping with the *fimW* promotor and/or regulatory elements (**Figure 2A**). These results suggest that silencing the expression of *fimW* leads to enrichment of STm at the ALI following repeated treatment with Sal4 IgA.

**Figure 2:**
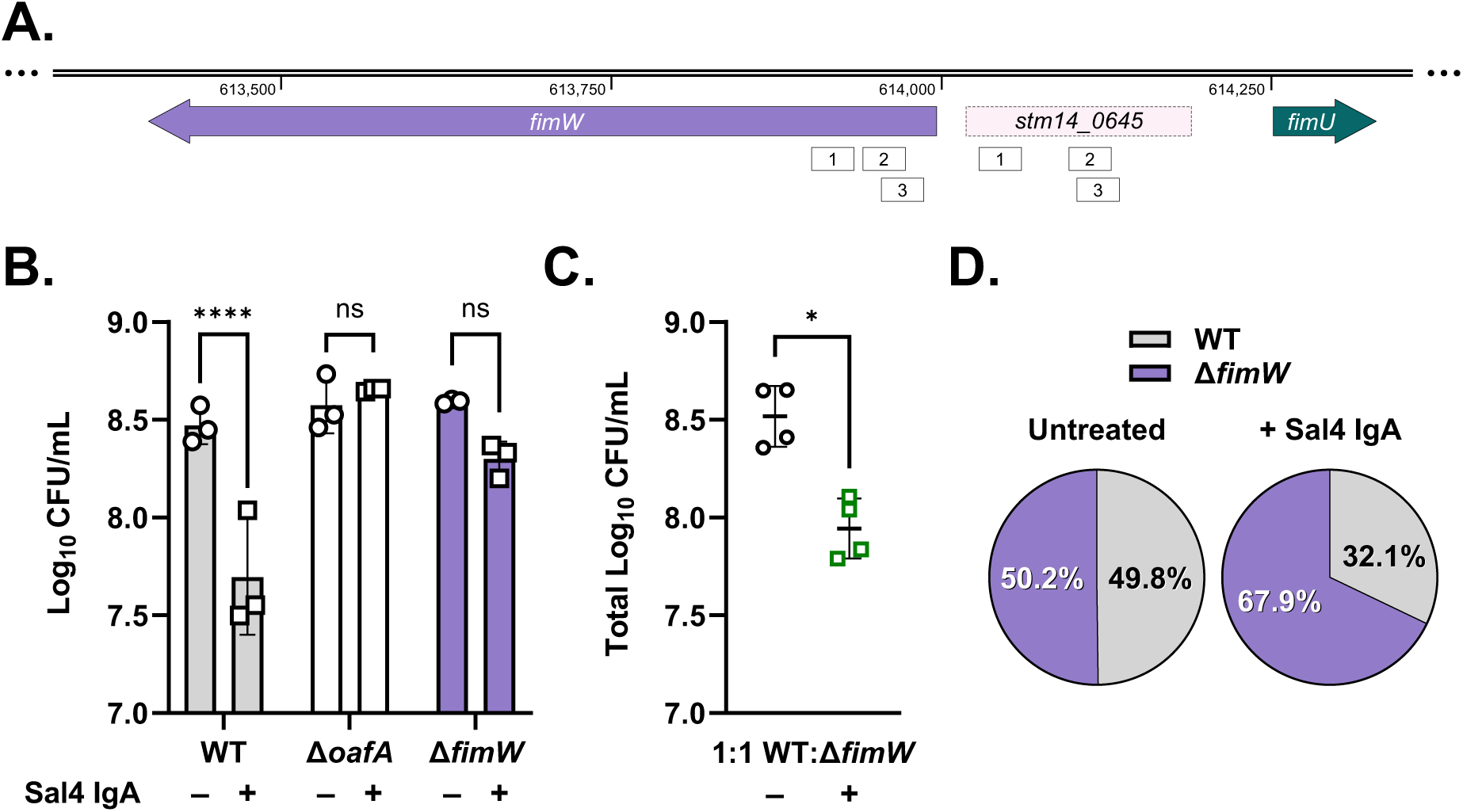
A STm *fimW* mutant evades Sal4-mediated agglutination *in vitro*. (A) Gene organization of *fimW* and neighboring genetic features, including the discontinued pseudogene *stm14_0645* and the tRNA-encoding gene *fimU*. White boxes represent enriched spacers (32 nucleotides) identified in the screen analysis that target complementary sequences within *fimW* and *stm14_0645*. For panels B and C, clearance of bacterial cells from the air-liquid interface of homogenous and mixed cultures of the indicated strains was quantified by plating colony forming units (CFUs) following 2 h of treatment with 15 μg/mL Sal4 IgA. (B) Data represents three biological replicates with error bars representing the standard deviation of the mean. Statistical significance was determined by ordinary two-way ANOVA with Šídák’s multiple comparison test. Asterisks (****) indicate p < 0.0001 and ns = not significant. (C) Data was obtained from four biological replicates with error bars representing the standard deviation of the mean. Statistical significance was determined by paired t-test (* indicates p < 0.05). (D) Percent composition of WT (SL174; *lacZ*^+^) and Δ*fimW* (SL164; *lacZ*^-^) recovered from the air-liquid interface as determined by blue-white screening of LB + X-gal plates. Values represent the average percentage of each strain from four biological replicates

We also reversed the parameters of the analysis to identify spacers that were de-enriched (i.e., hyper-agglutinators) at the ALI. A spacer was classified as de-enriched if its frequency in the untreated condition was >2-fold over that in the Sal4-treated condition. This analysis yielded 119 spacers, mapping to 106 unique genes and 8 promoters (**Dataset S4**). Three genes were targeted by 2 unique spacers each and two genes were targeted by 3 unique spacers each. Of these five genes, three encode proteins involved in flagellar complex assembly (*fliF, fliH,* and *fliN*). Seven additional genes each targeted by one spacer were also linked to flagellar biosynthesis and assembly. Selected genes of interest with their respective target spacers and log_2_-fold change values are shown in **Table 1**. These results suggest that down-regulation of flagellar-based motility renders cells more prone to agglutination upon Sal4 IgA treatment than motile counterparts. This is in contrast with the phenotype of non-motile mutants in monoculture, which do not agglutinate upon Sal4 treatment due to the lack of cell-cell collisions that drive classical agglutination (32). However, a 1:1 non-motile:motile mixture demonstrated equal agglutination of both strains due to cell-cell collisions caused by motile cells (32). From this, we speculate that the non-motile cells are entrapped into aggregates by collisions with motile cells and were disproportionally impacted by Sal4-mediated agglutination in the CRISPRi screen. As will be discussed later, motility may play opposing roles in Sal4 IgA-mediated agglutination depending on the proportion of motile cells in the immediate surroundings.

**Table 1:**
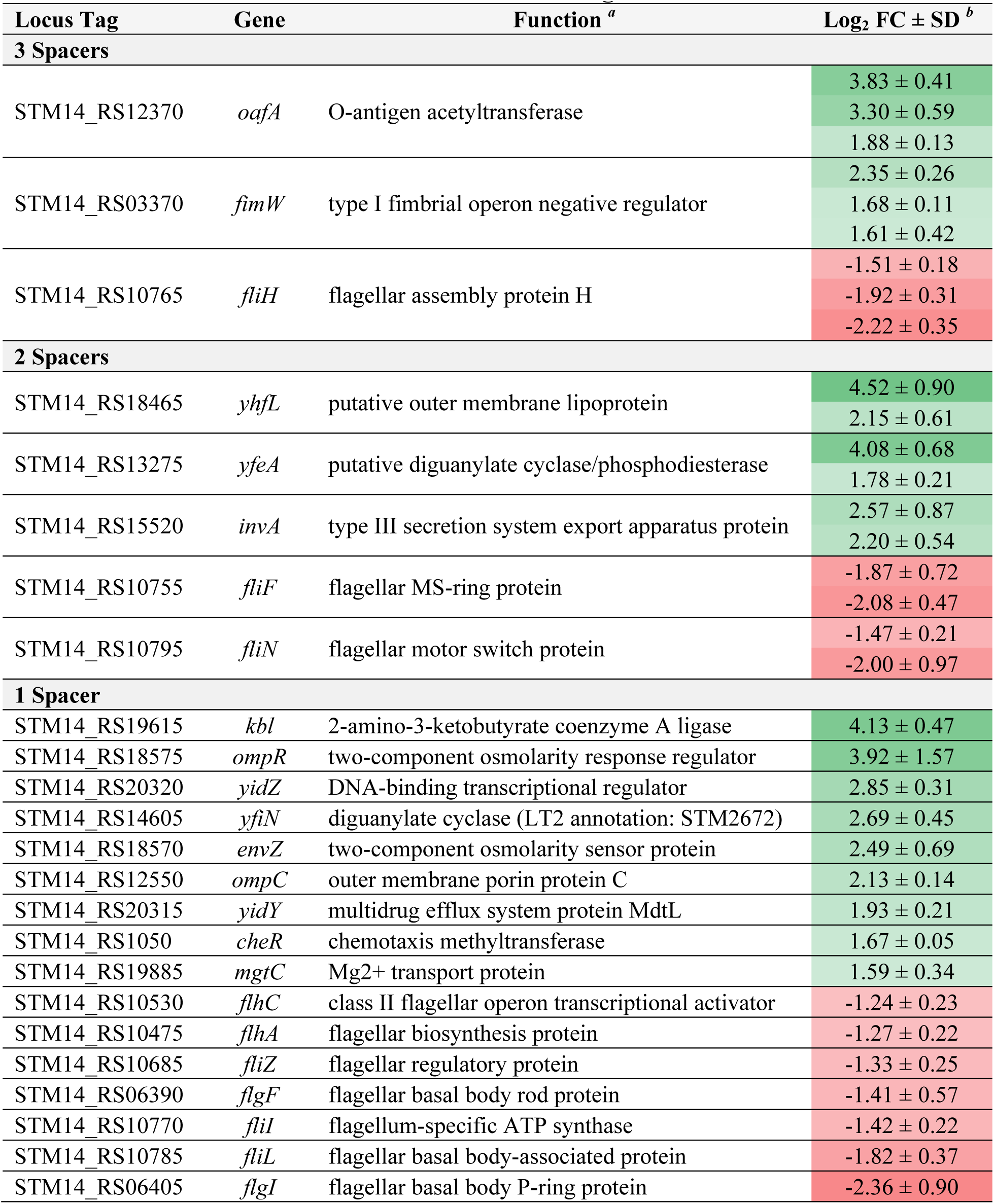
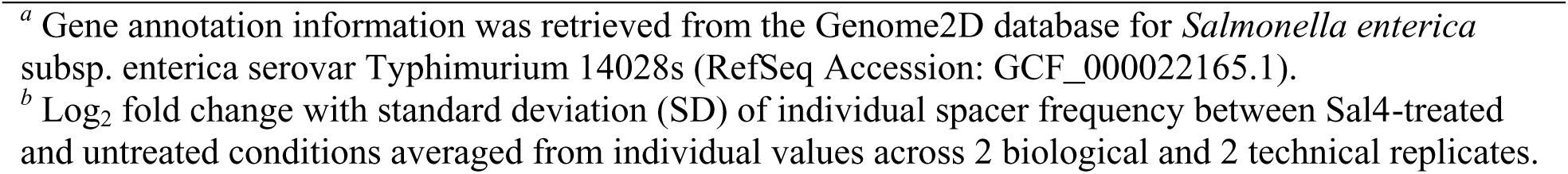
Select STm 14028s loci identified in the Sal4 IgA CRISPRi enrichment screen.

| Locus Tag | Gene | Function <sup>a</sup> | Log<sub>2</sub> FC ± SD <sup>b</sup> |
| --- | --- | --- | --- |
| 3 Spacers |  |  |  |
| STM14_RS12370 | oafA | O-antigen acetyltransferase | 3.83 ± 0.41 |
|  |  |  | 3.30 ± 0.59 |
|  |  |  | 1.88 ± 0.13 |
| STM14_RS03370 | fimW | type I fimbrial operon negative regulator | 2.35 ± 0.26 |
|  |  |  | 1.68 ± 0.11 |
|  |  |  | 1.61 ± 0.42 |
| STM14_RS10765 | fliH | flagellar assembly protein H | -1.51 ± 0.18 |
|  |  |  | -1.92 ± 0.31 |
|  |  |  | -2.22 ± 0.35 |
| 2 Spacers |  |  |  |
| STM14_RS18465 | yhfL | putative outer membrane lipoprotein | 4.52 ± 0.90 |
|  |  |  | 2.15 ± 0.61 |
| STM14_RS13275 | yfeA | putative diguanylate cyclase/phosphodiesterase | 4.08 ± 0.68 |
|  |  |  | 1.78 ± 0.21 |
| STM14_RS15520 | invA | type III secretion system export apparatus protein | 2.57 ± 0.87 |
|  |  |  | 2.20 ± 0.54 |
| STM14_RS10755 | fliF | flagellar MS-ring protein | -1.87 ± 0.72 |
|  |  |  | -2.08 ± 0.47 |
| STM14_RS10795 | fliN | flagellar motor switch protein | -1.47 ± 0.21 |
|  |  |  | -2.00 ± 0.97 |
| 1 Spacer |  |  |  |
| STM14_RS19615 | kbl | 2-amino-3-ketobutyrate coenzyme A ligase | 4.13 ± 0.47 |
| STM14_RS18575 | ompR | two-component osmolarity response regulator | 3.92 ± 1.57 |
| STM14_RS20320 | yidZ | DNA-binding transcriptional regulator | 2.85 ± 0.31 |
| STM14_RS14605 | yfiN | diguanylate cyclase (LT2 annotation: STM2672) | 2.69 ± 0.45 |
| STM14_RS18570 | envZ | two-component osmolarity sensor protein | 2.49 ± 0.69 |
| STM14_RS12550 | ompC | outer membrane porin protein C | 2.13 ± 0.14 |
| STM14_RS20315 | yidY | multidrug efflux system protein MdtL | 1.93 ± 0.21 |
| STM14_RS1050 | cheR | chemotaxis methyltransferase | 1.67 ± 0.05 |
| STM14_RS19885 | mgtC | Mg2+ transport protein | 1.59 ± 0.34 |
| STM14_RS10530 | flhC | class II flagellar operon transcriptional activator | -1.24 ± 0.23 |
| STM14_RS10475 | flhA | flagellar biosynthesis protein | -1.27 ± 0.22 |
| STM14_RS10685 | fliZ | flagellar regulatory protein | -1.33 ± 0.25 |
| STM14_RS06390 | flgF | flagellar basal body rod protein | -1.41 ± 0.57 |
| STM14_RS10770 | fliI | flagellum-specific ATP synthase | -1.42 ± 0.22 |
| STM14_RS10785 | fliL | flagellar basal body-associated protein | -1.82 ± 0.37 |
| STM14_RS06405 | flgI | flagellar basal body P-ring protein | -2.36 ± 0.90 |

<sup>a</sup> Gene annotation information was retrieved from the Genome2D database for *Salmonella enterica* subsp. enterica serovar Typhimurium 14028s (RefSeq Accession: GCF\_000022165.1).
<sup>b</sup> Log<sub>2</sub> fold change with standard deviation (SD) of individual spacer frequency between Sal4-treated and untreated conditions averaged from individual values across 2 biological and 2 technical replicates.

### A STm *fimW* null mutant evades Sal4-mediated agglutination in the SGA

To follow up on the results from the CRISPRi screen, we generated null mutants for 14 candidate genes with enriched spacers that encode products potentially involved in mediating agglutination in response to Sal4 treatment. Additionally, *flhC*, which encodes a master regulator of class II flagellar operons in STm, was chosen as a representative of the 10 flagellar-related genes associated with de-enriched spacers. We utilized the SGA to determine the extent of Sal4-mediated agglutination of each of the mutants relative to WT.

In the case of WT STm, the number of CFUs recovered following 2 h of Sal4 IgA treatment was reduced by ∼10-fold, as compared to untreated culture (8.45 ± 0.14 log_10_ versus 7.63 ± 0.27 log_10_) (**Table 2**). Conversely, the *oafA* mutant was recovered in equal numbers regardless of treatment (8.45 ± 0.19 log_10_ without versus 8.49 ± 0.18 log_10_ with Sal4 IgA), consistent with escape of antibody-mediated agglutination. In agreement with our previous study, the *flhC* mutant evaded Sal4-mediated agglutination as well (32). Surprisingly, among the 11 additional mutants we tested, only the *fimW* mutant remained enriched in the ALI following Sal4 IgA treatment, which echoed the phenotype of the *oafA* mutant (**Figure 2B**). The other 10 strains tested were as susceptible as WT to Sal4-mediated agglutination (**Table 2**).

**Table 2:**
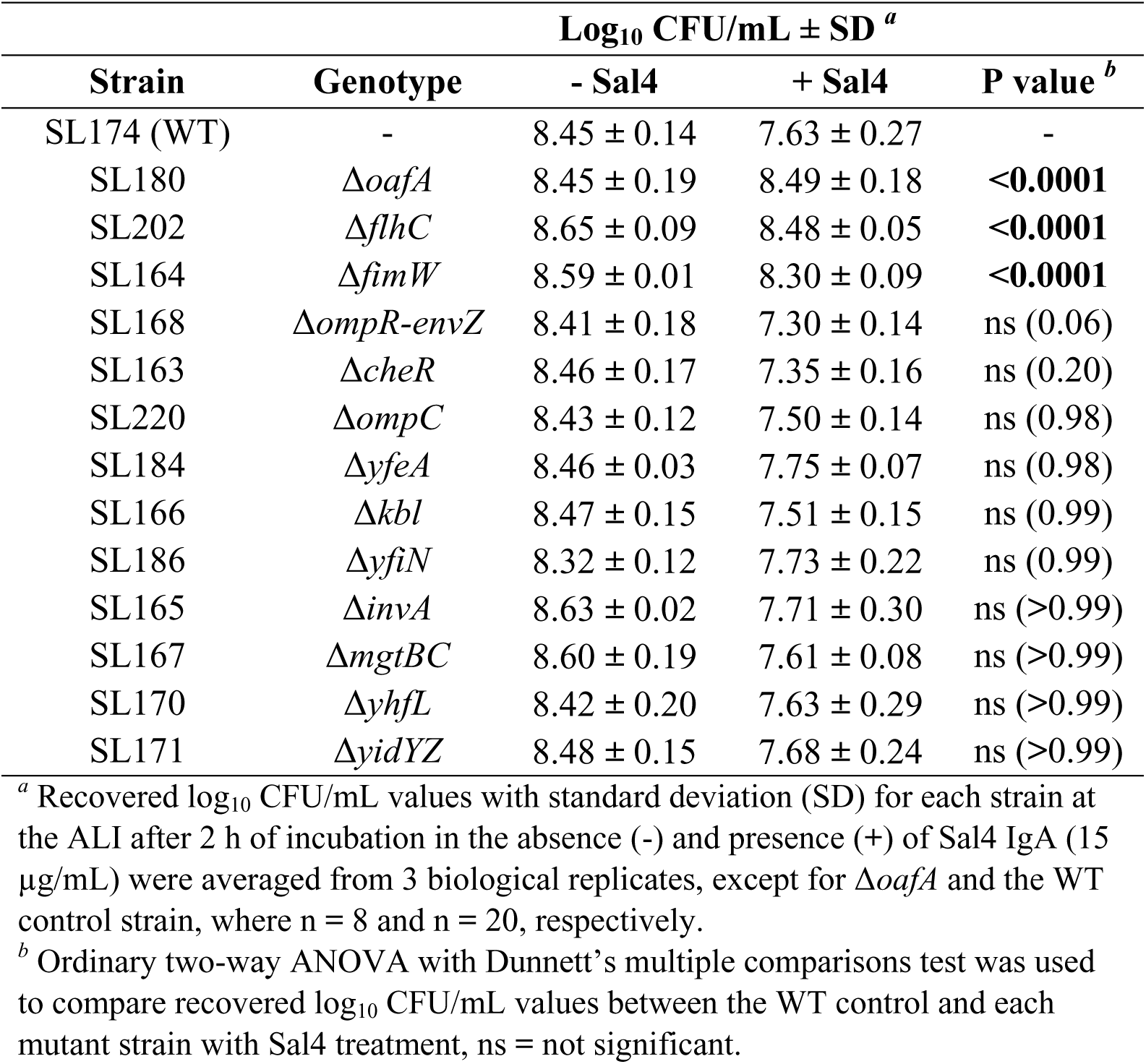
Sal4 IgA-mediated agglutination of STm mutants in the SGA.

The *fimW* gene encodes a negative regulator of Type I Fimbriae (T1F), extracellular mannose-sensitive filaments that facilitate host cell adhesion and pellicle formation under static culture conditions (36–40). Thus, we were particularly intrigued by a possible role for T1F in evasion of Sal4-mediated agglutination due to their reported association with pellicle formation (41). To confirm that loss of *fimW* does not simply result in inherent enrichment at the ALI, the *fimW* mutant (*lacZ^-^*) was mixed 1:1 with SL174 (WT *lacZ*^+^) in SGA in the absence or presence of Sal4 IgA. After 2 h, aliquots of the ALI were spotted on to LB-Xgal and the ratio of blue to white CFUs was compared. We observed a significant reduction in total CFUs recovered from the ALI upon Sal4 IgA treatment (**Figure 2C**). Blue-white screening of the colonies demonstrated that the *fimW* mutant and WT were present in nearly equal numbers in the absence of Sal4 IgA, while the *fimW* mutant was >two-fold more numerous that WT (67.9% to 32.1%) following Sal4 IgA treatment. These experiments validate the CRISPRi screen in that they confirm that loss of *fimW* expression results in evasion of Sal4 IgA-mediated agglutination, even within a mixed culture. In addition, whole cell ELISAs and dot blots were performed to rule out the possibly that the STm *fimW* mutant evades Sal4 IgA by altered O5-Ag expression. In both assays, Sal4 IgA reactivity with the *fimW* mutant was comparable to WT, demonstrating that the *fimW* mutation does not impact expression or accessibility of the O5 antigen (**Figure S2**).

### Phenotypic characterization and complementation of the STm *fimW* mutant

Previous studies have demonstrated that STm *fimW* null mutants are more fimbriate relative to WT STm due to overexpression of the *fim* operon (36). To confirm the hyperfimbriated phenotype of the *fimW* mutant, we developed a quantitative mannose-sensitive yeast agglutination (qMSYA) assay in a microplate format. In this assay, a change in optical density at 600 nm (ΔOD_600_) between the yeast culture alone compared to a culture mixed with STm is measured by spectrophotometry. STm strains with higher T1F expression are predicted to enhance yeast aggregation and decrease culture turbidity, resulting in a higher ΔOD_600_ value. As predicted, the *fimW* mutant produced visible yeast aggregates and had a significantly higher ΔOD_600_ value than that of WT STm (**Figure 3**). Moreover, the phenotype of the STm *fimW* mutant was similar to that of WT STm overexpressing *fimZ*, a positive transcriptional regulator of T1F, *in trans* (42). STm-induced agglutination of yeast was inhibited in the presence of 3% D-mannose, specifically implicating T1F in the agglutination phenotype. Both Sal4-mediated agglutination of the *fimW* mutant in the SGA and yeast agglutination caused by the *fimW* mutant in the qMYSA assay were restored to WT levels when the chromosomal *fimW* null mutation was complemented with a plasmid-encoded copy of *fimW* (pFimW) (**Figure 4AB**).

**Figure 3:**
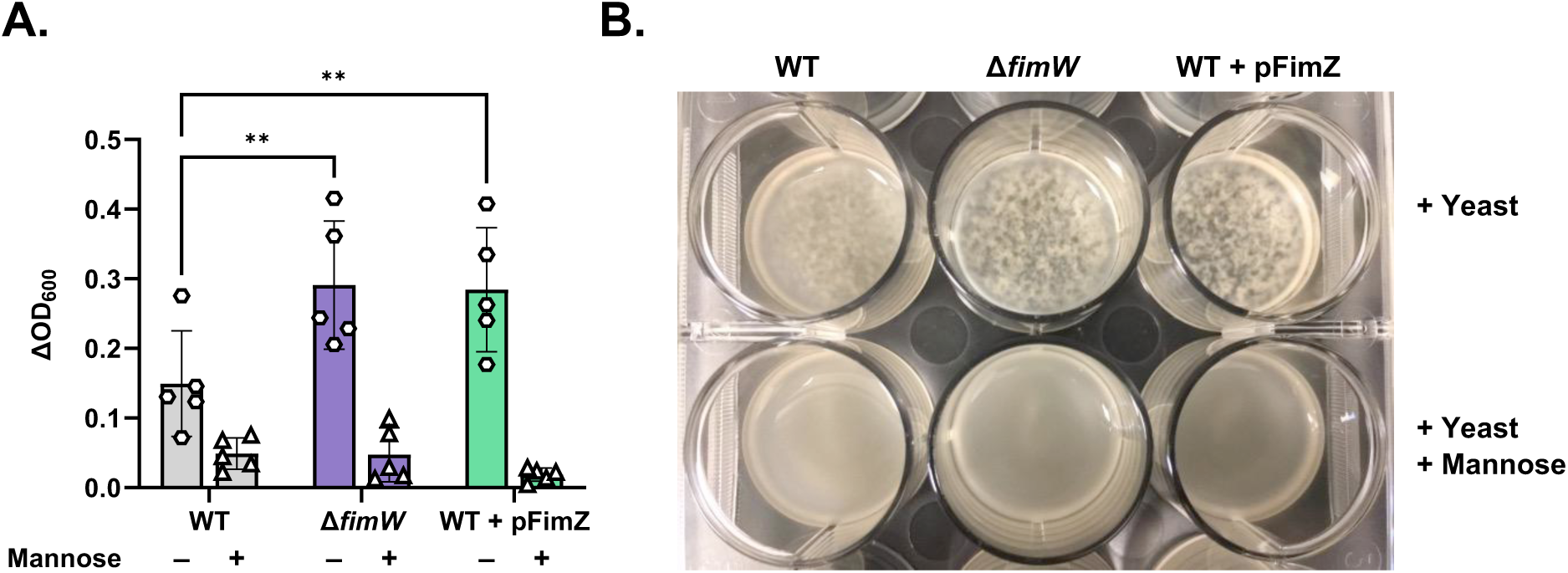
The *fimW* mutant efficiently agglutinates yeast in a mannose-sensitive manner. (A) Cultures of STm WT (SL174), Δ*fimW* (SL164), and WT + pFimZ (SL218; induced with 0.2% L-arabinose) were incubated statically for 48 h at 37 °C prior to centrifugation and resuspension in fresh LB. Cultures were mixed with yeast (final concentration: 10 mg/mL) either in the absence (hexagons) or presence of 3% (w/v) D-mannose (triangles) in a 12-well plate, as detailed in the Materials and Methods. The turbidity of the wells at 600 nm (OD_600_) was measured via spectrophotometry immediately after addition of yeast. Data was obtained from five biological replicates with error bars representing the standard deviation of the mean. Statistical significance was determined by two-way ANOVA followed by Tukey’s post hoc multiple comparisons test. Asterisks (**) indicate p < 0.01 and ns = not significant. (B) Representative image showing the extent of yeast agglutination when mixed with each STm strain in the absence and presence of mannose in the 12-well plate format.

**Figure 4:**
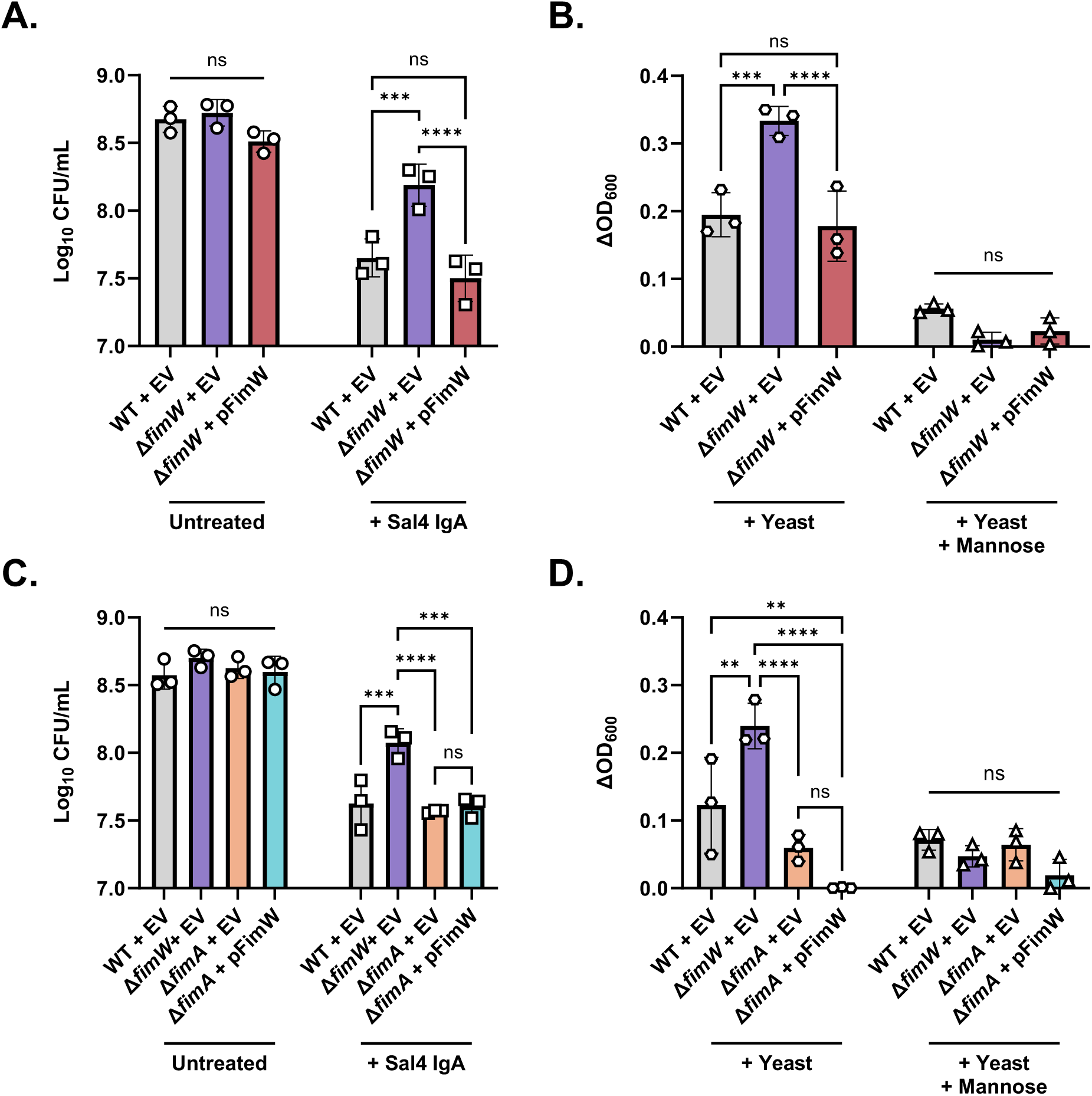
Overexpression of T1F reduces STm susceptibility to Sal4-mediated agglutination in the snow globe assay. (A) Recovered CFUs/mL of STm WT + pBAD24-EV (empty vector; EV), Δ*fimW* + EV, and Δ*fimW* + pBAD24-*fimW* (pFimW) cultures in the snow globe assay. (B) Quantification of mannose-sensitive yeast agglutination of the STm WT + EV, Δ*fimW* + EV, and Δ*fimW* + pFimW strains. (C) Recovered CFUs/mL of STm WT + EV, Δ*fimW* + EV, Δ*fimA* + EV, and Δ*fimA* + pFimW cultures in the snow globe assay. (D) Quantification of mannose-sensitive yeast agglutination of WT + EV, Δ*fimW* + EV, Δ*fimA* + EV, and Δ*fimA* + pFimW strains. For panels A and C, the indicated strains were grown to mid-log phase in the presence of 0.02% arabinose, washed in PBS, and either left untreated (circles) or treated with 15 μg/mL of Sal4 IgA (squares). After 2 h of treatment, the top of the supernatant was collected and plated on LB agar to measure CFUs. For panels B and D, the indicated strains were incubated statically for 48 h in LB containing 0.02% arabinose at 37 °C prior to centrifugation and resuspension in LB. Cultures were mixed with 10 mg/mL yeast in the presence (triangles) and absence (hexagons) of 3% mannose in a 12-well plate and the optical density of the wells at 600 nm (OD_600_) was measured via spectrophotometry. The strains used are SL257, SL253, SL255, SL289, and SL291. For all panels, data was obtained from three biological replicates with error bars representing the standard deviation of the mean. Statistical significance was determined by two-way ANOVA followed by Tukey’s post hoc multiple comparisons test. Asterisks (**, ***, ****) indicate p < 0.01, p < 0.001, and p < 0.0001, respectively, and ns = not significant.

### Direct implication of T1F in Sal4 IgA evasion

In addition to regulating T1F expression, FimZ and FimY also negatively regulate motility and invasion by down-regulating *flhDC* and upregulating *hilE*, respectively (43, 44). Therefore, we wanted to confirm that the non-agglutinating phenotype of the STm *fimW* mutant is solely due to the increased expression of T1F that occurs in the absence of FimW. To address this, we constructed a STm *fimA* mutant (which is afimbriate due to the lack of fimbrial subunit expression; Zeiner et al., 2012) then transformed it with either an empty pBAD24 vector (EV) or pFimW. The STm *fimA* mutant was as susceptible to Sal4 IgA-mediated agglutination as a WT + EV control strain in the SGA in both cases, and all three strains had significantly decreased recovered CFUs relative to Δ*fimW* + EV (**Figure 4C**). Additionally, the STm *fimA* mutants carrying either the pBAD24 empty vector or pFimW were defective in mannose-sensitive agglutination of yeast relative to the Δ*fimW* + EV and WT + EV strains (**Figure 4D**). Taken together, these results indicate that the reduced agglutination phenotype of the *fimW* mutant is directly due to the function of FimW as a negative regulator of T1F expression.

Previous studies have demonstrated that T1F expression in STm is induced in stationary growth conditions (38, 46). We reasoned that culturing STm under conditions conducive to TF1 expression would phenocopy a *fimW* null mutation and, at the same time, evade Sal4-mediated agglutination. WT STm was incubated with (T1F non-permissive) and without (T1F permissive) agitation in LB for 48 h at 37°C. The cultures were then subjected to the SGA and qMYSA. Cells derived from static culture evaded Sal4 IgA treatment in the SGA (as evidenced by higher CFUs in the ALI) and promoted yeast aggregation in the qMSYA as compared to cells derived from an analogous culture grown with agitation (**Figure S3**). These results are consistent with T1F playing a direct role in enabling STm to evade Sal4-mediated agglutination.

### The STm *fimW* mutant is vulnerable to additional Sal4 IgA-induced effects

Sal4 IgA has additional impacts on STm behavior and virulence beyond agglutination. Namely, Sal4 IgA is a potent inhibitor of STm flagellar-based motility in semi-solid agar, as well as an inhibitor of SPI-I T3SS activity and invasion of epithelial cells (19, 25). From this, we sought to determine whether the *fimW* mutant also evades motility arrest and epithelial cell invasion, in addition to Sal4-mediated agglutination. In a soft (0.3%) agar motility assay, the *fimW* mutant was indistinguishable from the WT control in both the absence and presence of Sal4 IgA, even at a lower concentration of antibody (5.0 µg/mL) (**Figures 5A** and S4). In contrast, motility of the *oafA* mutant was unaffected by Sal4 IgA (**Figure 5A**). This demonstrates that the hyperfimbriated phenotype does not enable STm to escape Sal4-mediated motility arrest.

**Figure 5:**
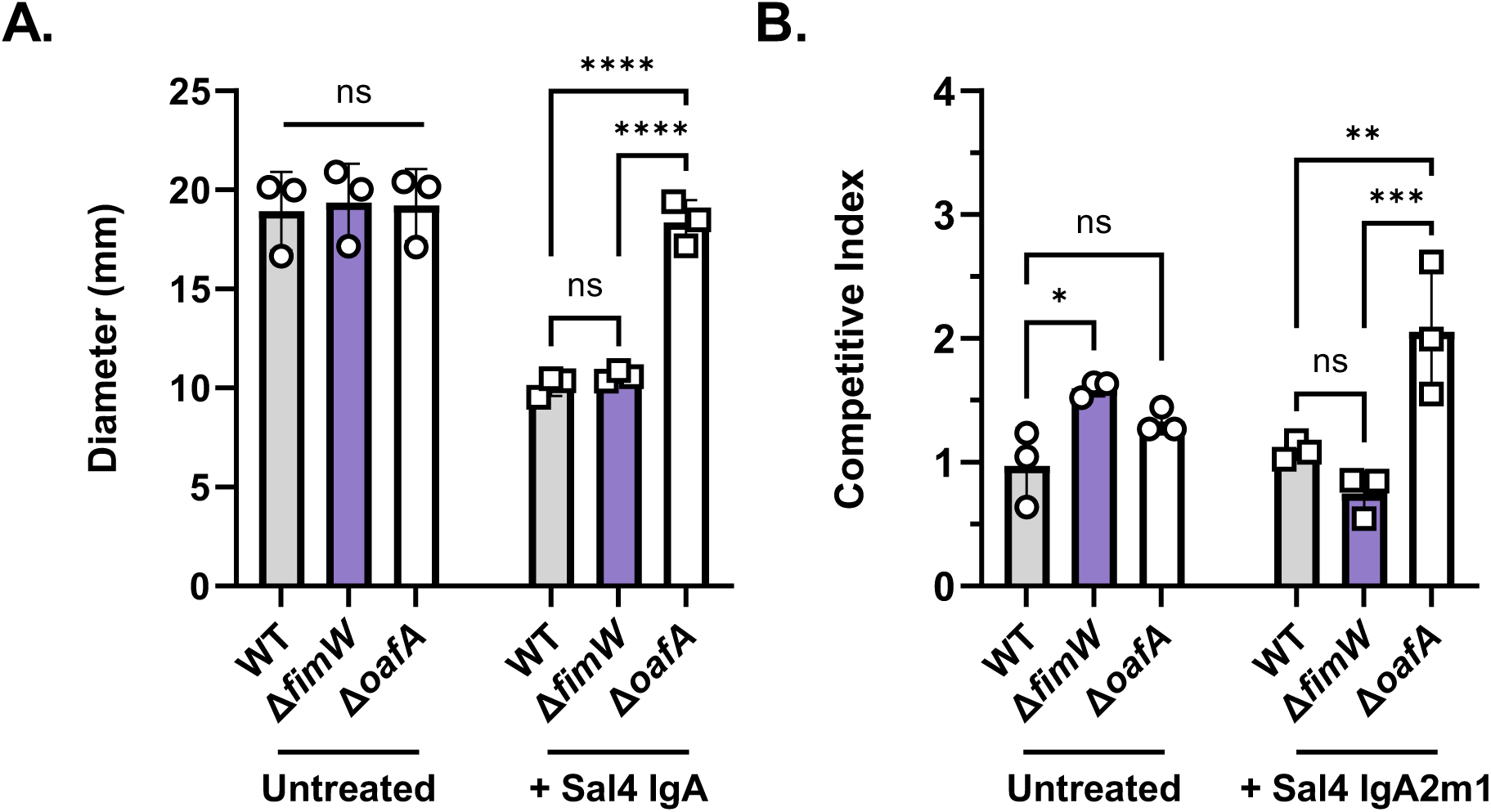
The *fimW* mutant is susceptible to additional Sal4-induced effects. (A) Plates of 0.3% LB agar with and without 5.0 μg/mL Sal4 IgA were stab inoculated with 1.0 μL of overnight cultures of WT (SL174), Δ*fimW* (SL164), and Δ*oafA* (SL180) and then incubated at 37°C for 4.5 h. Plates were imaged and the diameter (mm) of bacterial migration was measured using Fiji. Data represents three biological experiments each averaged from three technical replicates. Statistical significance was determined by two-way ANOVA followed by Tukey’s post hoc multiple comparisons test. Asterisks (****) indicate p < 0.0001 and ns = not significant. (B) WT (SL174; *lacZ*^+^) was mixed 1:1 with WT (SL239; *lacZ*^-^), Δ*fimW* (SL164; *lacZ*^-^), and Δ*oafA* (SL180; *lacZ*^-^) and incubated for 15 min with 15 μg/mL of purified recombinant Sal4 IgA2m1 before addition to HeLa cell monolayers in 96-well microtiter plates. Plates were centrifuged at 1000 x g for 10 minutes (rotating the plate after 5 min) to promote bacteria-cell contact. After 1 h of incubation at 37 °C, monolayers were treated with 100 µg/mL gentamicin and incubated again for 1 h to eliminate extracellular bacteria. Cells were washed and lysed with 1% Triton X-100 diluted in Ca^2+^ and Mg^2+^-free PBS and the resulting suspension was plated on LB containing X-gal to enable blue-white screening of CFUs. The competitive index [(%strain A output/%strain B output)/(%strain A input/%strain B input)] was calculated for each treatment group. Data represents three biological experiments each averaged from three technical replicates. Statistical significance was determined by two-way ANOVA followed by Tukey’s post hoc multiple comparisons test. Asterisks (*, **, ***) indicate p < 0.05, p < 0.01, and p < 0.001, respectively, and ns = not significant.

To evaluate the impact of Sal4 IgA on invasive fitness of the STm *fimW* mutant, we utilized a 1:1 mixed-strain competitive index (CI) invasion assay of human epithelial cells, as performed previously (25, 26, 28). In the absence of Sal4 IgA, the STm *fimW* mutant had a significantly higher CI value than that of the WT control, a result that is in line with the established role of T1F in promoting bacterial attachment to HeLa cells (**Figure 5B**) (40).

However, WT and the *fimW* mutant were similarly susceptible to Sal4 IgA, as evidenced by comparable CI values in the addition of antibody. By comparison, the STm *oafA* mutant had a >2-fold greater CI value than both WT and Δ*fimW* in the presence of Sal4 IgA (**Figure 5B**). These results demonstrate that the *fimW* mutant evades Sal4 IgA in the SGA but remains vulnerable to Sal4 IgA in motility and epithelial cell invasion assays. From this, we sought to determine the mechanism by which the STm *fimW* mutant selectively evades Sal4 IgA-mediated agglutination in the SGA.

### The STm *fimW* mutant forms a robust biofilm at the ALI following Sal4 treatment

T1F of STm have an established role in abiotic surface attachment and the formation of biofilms at the air-liquid interface (pellicles) in laboratory conditions (38, 39, 41, 47). We speculated that this function might account for the enrichment of the STm *fimW* mutant observed in the SGA, as we have reported that Sal4 IgA and IgG triggers extracellular matrix (ECM) production and biofilm formation of WT STm under certain conditions (20, 22). We utilized crystal violet (CV) staining to evaluate ECM secretion of the *fimW* mutant as compared to WT and Δ*oafA* controls. We also investigated the *flhC* mutant, as flagella have an established role in facilitating early stage biofilm formation (18, 48–51). Mid-log phase cultures of the four strains (WT, Δ*fimW,* Δ*flhC,* Δ*oafA*) were incubated in LB at 23°C with agitation for 1 h with and without Sal4 IgA (15 μg/mL) in a 12-well polystyrene microtiter plate and then stained with CV.

In the absence of Sal4 IgA, the four strains (WT, Δ*fimW*, Δ*oafA* and Δ*flhC*) produced minimal ECM, as evidenced by low levels of CV staining in the microtiter plates (**Figure 6**). The addition of Sal4 IgA did not cause ECM deposition by the WT, Δ*oafA* or Δ*flhC* strains, but did stimulate ECM production by the *fimW* mutant, as evidenced by the >3-fold increase in Abs_570_, correlating with enhanced CV staining. This is in agreement with previous studies that have noted Sal4 IgA does not trigger biofilm formation in WT STm at room temperature (20). Thus, the *fimW* mutant is prone to ECM production under non-inducing conditions for WT STm with Sal4 IgA treatment.

**Figure 6:**
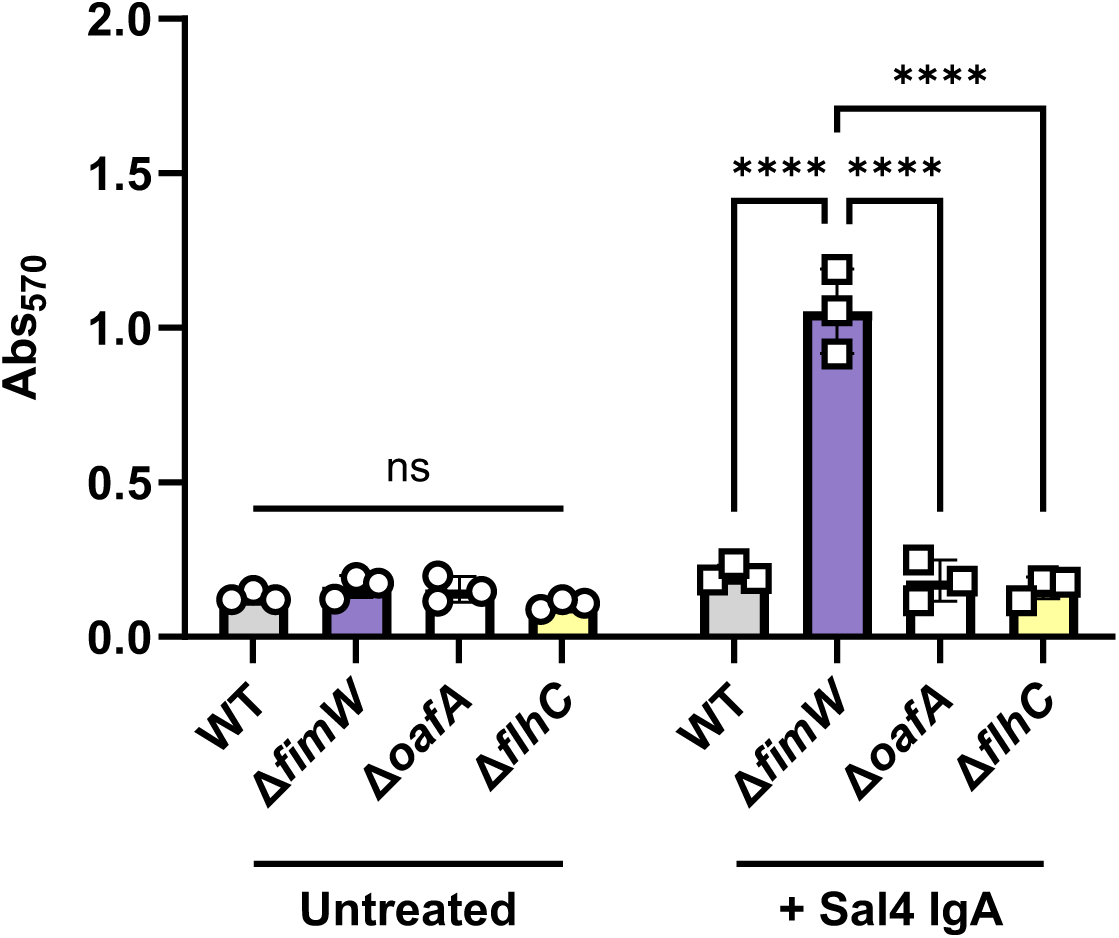
Overexpression of T1F enhances Sal4 IgA-mediated biofilm production on polystyrene. Mid-log phase cultures of the indicated strains were washed with LB, standardized to an OD_600_ value of 1.0, and transferred to a 12-well polystyrene tissue culture plate. Cultures were left untreated (circles) or treated with 15 μg/mL Sal4 IgA (squares) at 23°C with shaking (200 rpm) for 1 h. The culture media was aspirated, and the biofilms were heat-fixed at 60°C for 1 h. Biofilms were stained with 0.1% crystal violet and washed with distilled water for 5 min, then the CV stain was solubilized with 30% acetic acid for 5 min. CV stain absorbance was quantified at Abs_570_. Data represents three biological experiments and error bars represent standard deviation of the mean. Statistical significance was determined by two-way ANOVA followed by Tukey’s post hoc multiple comparisons test. Asterisks (****) indicate p < 0.0001 and ns = not significant.

We next examined ECM production of the same four strains when cultured in borosilicate glass tubes with shaking in the presence of purified Sal4 IgG. In the absence of antibody treatment, the STm WT, Δ*fimW,* Δ*oafA* and Δ*flhC* strains again showed low levels of CV staining. In contrast, after 1 h of treatment with Sal4 IgG, the WT, Δ*fimW* and Δ*flhC* strains formed visible rings of ECM-like material at the ALI, albeit with varying degrees of thickness and durability (**Figure 7A**). The WT strain formed a relatively uniform band that was readily stained with CV, while the Δ*flhC* strain formed a thinner band with interspersed gaps that was loosely adhered to the glass surface, resulting in partial removal during processing. In contrast, the *fimW* mutant formed a thick band with small projections emanating from the ring, which resulted in intense CV staining. When the CV was solubilized and quantitated by spectrophotometry, both WT and STm Δ*fimW* had significantly higher Abs_570_ values than those Δ*oafA* and Δ*flhC* (**Figure 7B**). Additionally, STm Δ*fimW* demonstrated greater CV staining than the WT, likely due to enhanced adhesion to the glass mediated by T1F. Collectively, these results provide an explanation for the observed enrichment of *fimW*-targeting spacers in the CRISPRi screen and the evasion phenotype of the *fimW* mutant at the ALI after 2 h of Sal4 IgA treatment in the SGA.

**Figure 7:**
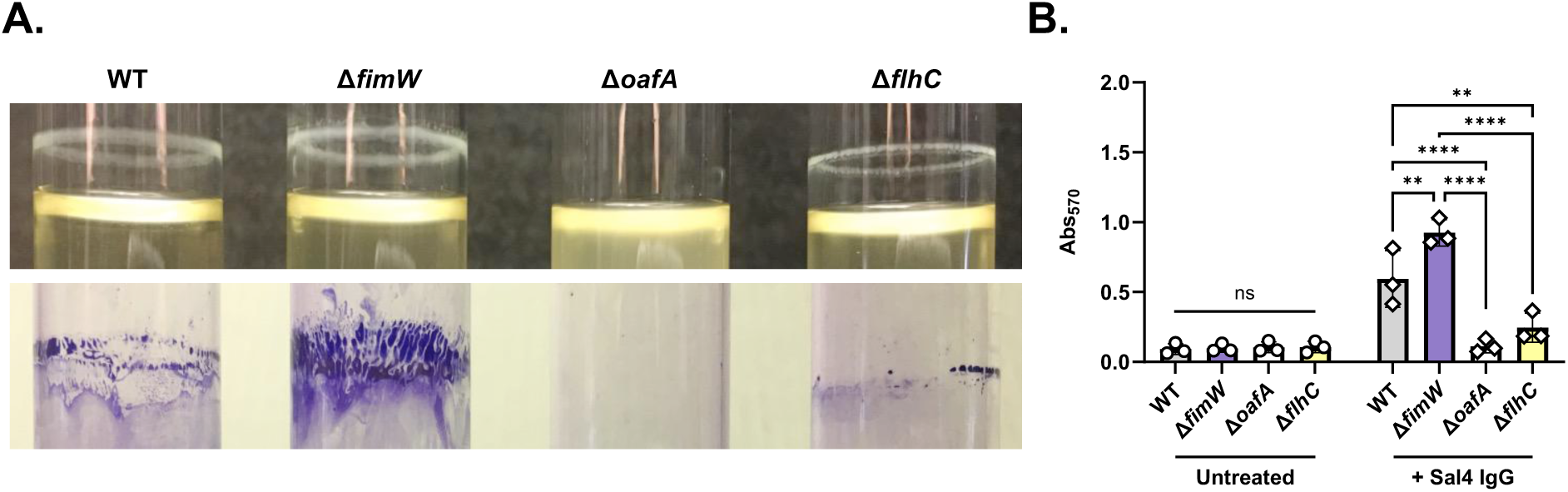
CV staining of agglutination-evading mutants following Sal4 IgG treatment. Mid-log phase cultures of WT (SL174), Δ*fimW* (SL164), Δ*oafA* (SL180), and Δ*flhC* (SL202) were washed with LB, standardized to an OD600 value of 1.0 and transferred to borosilicate glass tubes. Cultures were mixed 1:1 with LB with (diamonds) and without (circles) 30 μg/mL Sal4 IgG and then incubated at 23°C with shaking (200 rpm) for 1 h (final concentration of 15 μg/mL Sal4). The culture media was aspirated, and the plate/tubes were heat-fixed at 60°C for 1 h. Biofilms were stained with 0.1% crystal violet and washed with distilled water for 5 min each, then the CV stain was solubilized with 30% acetic acid for 5 min. Dissolved CV was transferred to a 96-well plate prior to measurement of absorbance at 570 nm (Abs_570_). (A) Representative images of the indicated strains 1 h p.t. (top) and with remaining CV stain after the final wash step (bottom). (B) Data represents three biological experiments and error bars represent standard deviation of the mean. Statistical significance was determined by two-way ANOVA followed by Tukey’s post hoc multiple comparisons test. Asterisks (**, ****) indicate p < 0.01 and p < 0.0001, respectively, and ns = not significant.

## DISCUSSION

Secretory IgA plays a central role in protecting the intestinal epithelium from bacterial pathogens like STm. At low bacterial cell density, IgA cross-links daughter cells and prevents their separation through a phenomenon known as enchained growth (17). At higher cell densities, IgA mediates “classical” agglutination of pathogens, which entails cross-linking of neighboring cells via antibody-mediated intercellular bridges and the eventual formation of macroscopic aggregates that share hallmarks associated with bacterial biofilms (22, 52, 53). In many ways, classical agglutination serves as a paradigm for understanding how an external cue (i.e., IgA) drives conversion of bacteria from a planktonic, infectious state to an aggregated, non-infectious one. In the case of STm, understanding the molecular events underlying this transition may have implications for mucosal vaccine development and oral immunotherapies.

In this report, we performed a STm genome-wide CRISPRi screen to identify genes whose expression influenced dynamics of Sal4 IgA-mediated agglutination at the ALI. The screened uncovered *fimW*, a negative regulator of T1F, which are known to be involved in pellicle formation and ALI interactions (38, 39, 41). We generated a STm *fimW* null mutant and confirmed that the strain was phenotypically hyperfimbriated with a fully quantitative yeast agglutination assay. More importantly, the *fimW* mutant strain evaded Sal4 IgA-mediated agglutination in the SGA, even though it was still susceptible to Sal4 IgA in both the motility and HeLa cell invasion assays. Further examination of the ALI revealed that the *fimW* mutant was prone to enhanced biofilm formation at the ALI. Conversely, flagella biosynthesis and assembly genes, including the master transcription factor gene *flhC*, were significantly de-enriched in the CRISPRi screen, implicating expression of the flagella in Sal4-mediated biofilm formation.

Taken together, we propose a multistep model in which Sal4 IgA triggers STm to undergo a planktonic-to-sessile transition that culminates multicellular aggregation (**Figure 8**). The requisite first step is the recognition of the O5-antigen by Sal4 IgA, which may mimic cell surface contact and/or a form of outer membrane stress (19, 20, 24–26, 31). The observed enrichment of spacers targeting *oafA* in the CRISPRi screen confirms the importance of engaging the O-antigen in initiating downstream events. In step 2, flagella-based motility promotes physical contact between STm and inert surfaces. At this stage flagella may also play a role in surface sensing and activating signal transduction pathways, possibly involving second messengers like c-di-GMP (54, 55). At the same time, free-floating bacterial aggregates are formed as the result of cell-cell collisions (22, 32, 56, 57). In step 3, adhesins facilitate bacterial attachment to abiotic materials and the ALI. In the case of the SGA, we propose that the T1F serves this function, possibly due to enhanced hydrophobic interactions with these surfaces (47, 58, 59). In retrospect, the experimental conditions of the CRISPRi screen (i.e., prolonged static incubation in nutrient-poor media) were ideal for enriching spacers targeting *fimW,* as T1F have an established role in pellicle formation (38, 41). That said, it is worth noting that among the 13 types of fimbriae encoded on the STm genome, T1F appear to be the primary mediators of adherence to abiotic surfaces in laboratory conditions (60). Finally, in step 4, we propose that STm biomass increases as the result of the capture and retention of Sal4-coated single cells and/or multicellular aggregates. While this four step model is based principally on the results of the SGA, studies examining IgA-mediated agglutination of STm in mouse models of intestinal infection corroborate the sequence of events, including the formation of large and densely packed biofilm-like bacterial aggregates in the small intestine (22).

**Figure 8:**
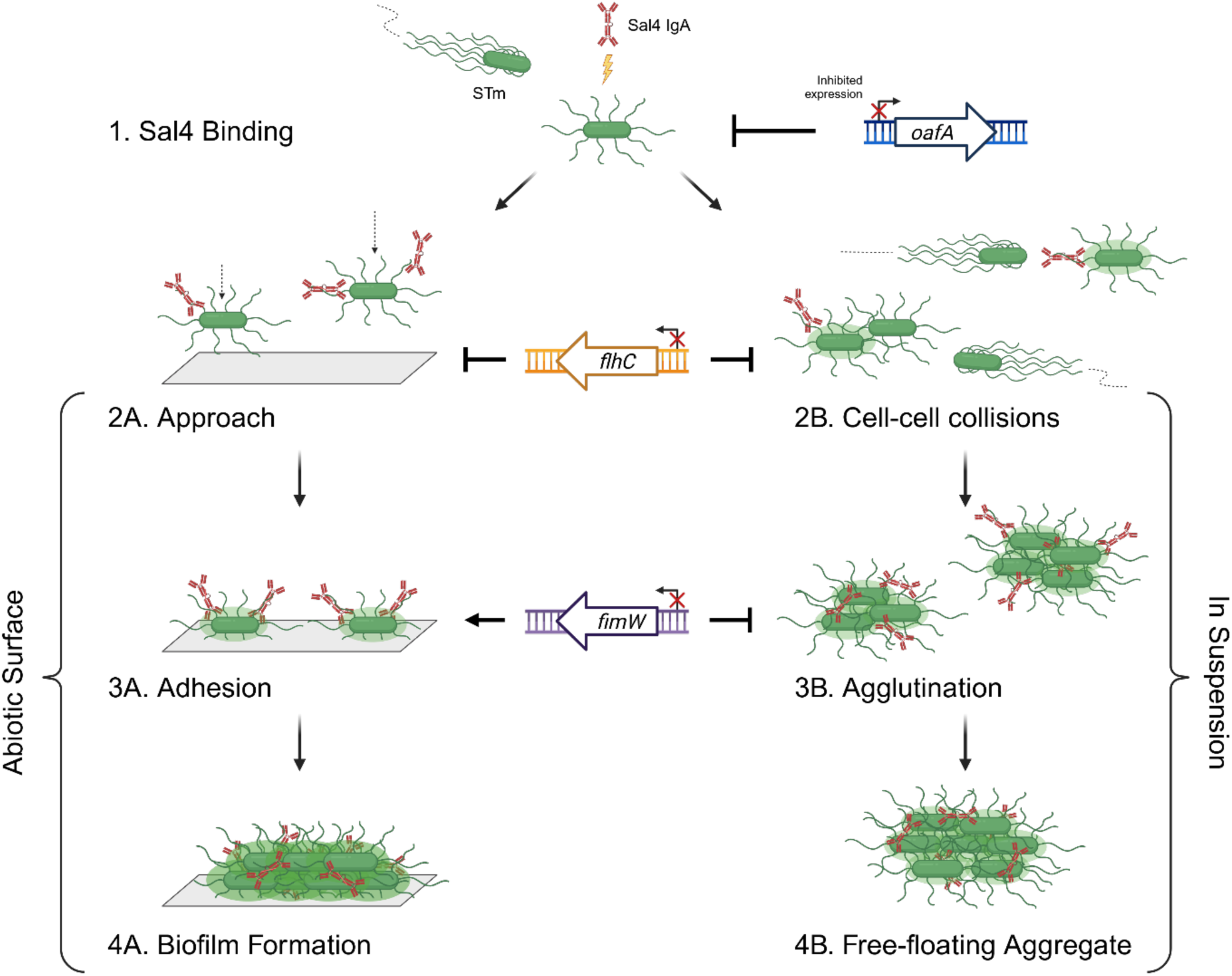
Proposed model of Sal4-induced impacts on STm *in vitro*. Sal4 IgA binding to the O5 antigen (dependent on expression of *oafA*) induces a form of outer membrane stress that signals STm to transition from a planktonic to sessile state, either proximal to an abiotic surface or in suspension (step 1). Using flagellar-based motility, the bacteria swim towards an abiotic surface (step 2A) and collide into each other (step 2B). Fimbriae and Sal4-induced EPS production facilitate adhesion to the surface (step 3A). Hyperfimbriation, occurring through inhibition of *fimW*, enhances this step and reduces the extent of agglutination of STm in suspension (step 3B). Free-floating aggregates and surface-adhered microcolonies both bound by Sal4 IgA continue to produce EPS, resulting in the formation of biofilms (step 4AB).

Based on the results of this study, the role of flagellar-based motility in Sal4-mediated agglutination is multifaceted. A 1:1 mixed culture of motile and non-motile strains of STm resulted in equal agglutination of both strains after 2 h of Sal4 IgA treatment, indicating that non-motile mutants are just as vulnerable to agglutination as a WT motile strain, as long as there is a nucleating factor present to initiate aggregate formation (32). From this observation, we proposed a bystander catch model, where non-motile bacteria that do not agglutinate on their own are entrapped in aggregates from collisions with their motile counterparts. In the pooled CRISPRi mutant library, cells with spacers targeting flagella-related genes would be a fraction of the rest of the culture containing unrelated spacers, indicating the former faces increased susceptibility to Sal4-mediated agglutination in a biased population. The *flhC* mutant formed a relatively fragile biofilm, aligning with the established role of flagella in surface approach and providing a scaffold for other cells to adhere to during the early stages of biofilm formation (18, 48–51). Along with the observation that the *fimW* mutant forms a robust biofilm, this suggests that the subpopulation of STm at the ALI are prone to forming a biofilm, while those that are de-enriched represent biofilm-deficient cells. Future studies are necessary to determine the biofilm formation capacity and composition of the other null mutant strains generated in this study.

We have repeatedly noted the parallels between antibody-mediated bacterial agglutination and biofilm formation, including a possible role for the secondary messenger, cyclic-di-GMP (20, 32, 61, 62). For example, Amarasinghe *et al.* implicated YeaJ, an inner membrane-localized diguanylate cyclase (DGC), in regulating EPS production following Sal4 exposure (20). Thus, we were not surprised to enrich for spacers targeting several cyclic-di-GMP metabolizing enzymes (CMEs) in the CRISPRi screen. While those specific CMEs did not appear to contribute to Sal4 IgA-mediated agglutination in the SGA in follow-up studies, nor did we identify *yeaJ* in our screen, further work is necessary before ruling out the involvement of c-di-GMP in the response to Sal4. Of particular interest is a putative phosphodiesterase-encoding gene (*stm14_0643* in STm 14028s; *stm0551* in STm LT2) located between *fimW* and *fimY* (63). A *stm0551* mutant was shown to produce T1F on solid agar and this correlated with increased expression of *fimA* and *fimZ* (63). Furthermore, PDE activity of the purified protein was confirmed in this study, suggesting that c-di-GMP is directly involved in regulation of T1F (63). While we did not identify spacers targeting *stm14_0643* in the CRISPRi screen, we are actively investigating its possible role in facilitating Sal4-induced biofilm formation in STm (S. Lindberg and N. Mantis, unpublished results).

Taken together, the results from this study implicate T1F in promoting STm survival in the external environment, in addition to their well-established function as a virulence factor.

Overexpression of T1F appears to enhance biofilm formation of STm that is induced by Sal4 IgA treatment, while a *fimW* mutant is still vulnerable to other Sal4-induced effects like motility arrest and inhibition of invasion *in vitro.* The results from this study indicate that the lack of O5 antigen expression may be the only one true method of Sal4 escape, as complete resistance to Sal4 has been exclusively observed with an *oafA* mutation thus far. Thus, this study demonstrates the effectiveness of Sal4 as a neutralizing agent against STm, as long as its epitope is present.

The ability of Sal4 to induce biofilm formation of STm has implications for eventual shedding and transmission of bacterial aggregates following clearance from the intestinal lumen. Further studies are necessary to determine the underlying mechanisms of Sal4 IgA-induced biofilm formation of STm and better understand how SIgA mediates protection at mucosal surfaces.

## MATERIALS AND METHODS

### Bacterial strains and growth conditions

Bacterial strains and plasmids used in this study are listed in Supplementary Tables 1 and 2, respectively*. Salmonella enterica* serovar Typhimurium (STm) strain 14028s was obtained from the American Type Culture Collection (ATCC, Manassas, VA). Bacteria were cultured as described (32). Cultures of 5 mL Luria-Bertani (LB) medium were inoculated with a single isolated colony from a freshly streaked LB agar plate and grown overnight (∼16 h) at 37°C with aeration (225 rpm) in a MaxQ 4000 benchtop incubator (ThermoFisher Scientific, Waltham, MA). Overnight cultures were subcultured 1:50 into LB broth and grown to mid-log phase (OD_600_ of ∼0.7) prior to experimentation. Optical density at 600 nm (OD_600_) was monitored using a GENESYS 10S UV-Visible spectrophotometer (ThermoFisher Scientific). When appropriate, growth media was supplemented with kanamycin (50 µg/mL), carbenicillin (100 µg/mL), and/or 5-bromo-4-chloro-3-indolylbeta-D-galacto-pyranoside (X-gal; 40 µg/mL). L-arabinose was added to a final concentration of 0.2% or 0.02% to induce pBAD promoters, as indicated in the text (SigmaAldrich, St. Louis, MO). Strains carrying a temperature-sensitive plasmid were maintained at 30°C (64, 65).

### Generation of STm mutants

Mutant strains were generated as described (32). Lambda Red recombination was utilized to generate STm strains with the target gene replaced by an antibiotic resistance cassette (64). The Kanamycin resistance cassette from pKD13 (64) was amplified using flanking primers and the resulting PCR products were purified using a DNA Clean & Concentrator-5 kit according to the manufacturer’s instructions (Zymo Research, Irvine, CA). STm 14028s carrying the pKD46 plasmid (64) (strain SL094) was grown in the presence of 0.4% arabinose (to induce expression of the Lambda Red recombinase genes encoded by pKD46), washed four times with 10% glycerol, transformed with the concentrated PCR product via electroporation, then recovered in LB for 1 h at 30°C with aeration. Recovered cells were plated onto LB agar containing kanamycin and incubated overnight at 37°C. Recombinants were verified using colony PCR for antibiotic resistance cassette insertion at the correct genomic location. Finally, to generate a constitutively *kanR-*expressing strain of STm 14028s, WT STm strain harboring pTn7 (65) (SL172) was cultured to mid-log-phase in LB supplemented with carbenicillin and 0.4% arabinose and then electrotransformed with the Tn7 transposition plasmid pUC18-R6k-mtn7-kanR (66). Transformants were plated onto LB agar containing kanamycin and incubated overnight at 37°C. The pKD46 and pTn7 plasmids were cured by incubating the newly generated mutant strains for 4 h at 42°C with aeration in the absence of antibiotic selection. Oligonucleotide primers used to construct plasmids and STm mutant strains are listed in Supplementary Table 3.

### Construction of pFimW and pFimZ plasmids

Plasmids overexpressing *fimW* and *fimZ* were prepared essentially as described (20). The coding sequences of *fimW* and *fimZ* from the STm 14028s genome were amplified via PCR with *fimW*- and *fimZ*-specific primers (**Table S3**) engineered with restriction sites for XbaI and HindIII. The resulting gene fragments were cloned into a XbaI/HindIII-digested pBAD24 vector (67) using T4 DNA ligase (NEB, Ipswich, MA).

The resulting plasmids were cloned into competent *E. coli* DH5α F’*I^q^* cells and transformants were selected on LB agar plates containing carbenicillin (NEB). The DNA sequence of the plasmid inserts for each transformant was verified before and after transformation into STm by electroporation. To generate STm strains carrying pFimW and pFimZ, the plasmids were purified from *E. coli* DH5α F’*I^q^* using a QIAprep Spin Miniprep Kit (QIAGEN, Germantown, MD).

### Monoclonal antibodies and hybridomas

The B cell hybridoma cell line secreting monoclonal polymeric Sal4 IgA was maintained as described (23, 25). Recombinant Sal4 IgA2m1 was provided by Moderna, Inc. (Cambridge, MA) and chimeric Sal4 IgG_1_ was provided by MappBio, Inc (San Diego, CA) (26, 28).

### Snow globe assay (SGA)

The snow globe assay was performed as described (**32**). In brief, overnight cultures (or 48 h cultures for Figure S2) of STm were subcultured 1:50 in fresh media and grown to mid-log phase (OD_600_ = ∼0.7) with aeration at 37°C. The cultures were pelleted by centrifugation in a Heraeus Multifuge X1R Centrifuge (Thermo Fisher) at 6000 x g for 4 min and resuspended in sterile PBS (pH 7.4). The centrifugation and resuspension in PBS steps were repeated for a total of two washes, then the OD_600_ of the cultures were adjusted to a value of 1.0 using PBS. Cultures were transferred to a borosilicate glass tube (16 x 25 mm) and either left untreated or treated with 15 μg/mL Sal4 IgA in a final volume of 5 mL. Bacterial agglutination was recorded using an iPhone 6s (Apple, Cupertino, CA) with the ‘TimestampCamera’ application. To obtain colony forming units (CFUs), 100 µL was taken from the very top of the culture tube at the ALI, serial diluted fivefold in sterile PBS pH 7.4, and then 100 µL from two consecutive dilutions was plated on LB agar and spread using glass plating beads. Plates were incubated at 37°C overnight in a Heratherm IMH60 incubator (ThermoFisher Scientific) overnight and counted the following day using an eCount Colony Counter (Heathrow Scientific, Vernon Hills, IL). CFU counts from the dilution plates were averaged to calculate a CFU/mL value for each condition, with plates having more than 300 or less than 30 colonies excluded from the final dataset.

### Mutant enrichment SGA

STm cultures were grown from overnight cultures to mid-log phase and prepared for experimentation as described above. An *oafA* mutant strain (SL180) was spiked into a WT culture (SL174; *lacZ^+^*) at a ratio of 1:6000 (approximately 4.6 E+05:2.7 E+09 CFUs, respectively) and then mixed thoroughly by gentle pipetting. The mixed cultures were then left untreated or treated with 15 μg/mL Sal4 IgA in a final volume of 7.5 mL. After incubation at room temperature for 5 hours, both untreated and treated cultures were passaged by transferring 10 μL from the top of the culture tube into 5 mL fresh media and incubated at 37°C and 220 rpm overnight. In addition, 100 μL was taken from the ALI, serial diluted fivefold in sterile PBS (pH 7.4), and plated on LB + Kan + X-gal. Plates were incubated overnight at 37°C and CFUs were counted the following day. This procedure was repeated for subsequent rounds of treatment.

### CRISPRi macroagglutination screen

Prior to experimentation, the previously designed CRISPRi library strain (SL061) was grown up from a frozen aliquot in 50 mL LB supplemented with carbenicillin overnight at 37°C with aeration. The following morning, 50% glycerol was added to the culture (final concentration of 10%) and incubated for 5 minutes with aeration to mix thoroughly. Aliquots of 250 μL were added to presterilized 1.5 mL microcentrifuge tubes and stored at -80°C until use. To perform the screen, aliquots of the library strain were thawed and grown to mid-log phase in LB supplemented with carbenicillin at 37°C with aeration. In addition, an overnight culture of SL118 (*zjg8101::kan oafA126::Tn10d-Tc fkpA-lacZ* + pBAD24-EV; used as a visual indicator of enrichment) was subcultured 1:50 in LB + Carb and grown to mid-log phase at 37°C with aeration. Both cultures were centrifuged, washed twice in PBS pH 7.4, and standardized to an OD_600_ of 1.0. The SL118 culture was mixed with that of SL64 at a ratio of 1:10,000 and then either left untreated or treated with 15 µg/mL Sal4 IgA in a final volume of 7.5 mL. After 5 h of treatment at room temperature, 200 µL was taken from the ALI to serial dilute and plate for CFUs and 15 µL was used to passage the cultures overnight. The next day, overnight cultures were diluted 1:50 in LB + Carb and grown to mid-log phase prior to preparation, treatment, plating, and passaging as described above. The following day, plasmids containing spacer sequences were isolated from 3 mL of overnight culture from each sample with a QIAprep Spin Miniprep Kit (QIAGEN) according to the manufacturer’s instructions. The concentration of purified DNA was quantified by Nanodrop (ThermoFisher Scientific). The region of the plasmid encoding the spacer was amplified with a low (∼10X) amplification cycle PCR program and PCR products were resolved on an 8% polyacrylamide gel. The resulting DNA pool was purified using AMPure XP reagent (at a ratio of 0.8 beads/sample volume) according to the manufacturer’s instructions (Beckman Coulter, Indianapolis, IN). Final DNA concentration was quantified using a Qubit 4.0 fluorometer and dsDNA Broad-Range Assay Kit (ThermoFisher Scientific). Samples were then submitted to the AGT Core at the Wadsworth Center and run on an Illumina NextSeq 500 instrument (San Diego, CA). A custom Python script was utilized to extract the number of reads for each spacer from the NGS .fastq data files (Dataset S2) and further analyzed using a custom R script (Supplementary Script Figure/File 1). The screen was performed twice, each with two technical replicates.

### Quantitative mannose-sensitive yeast agglutination assay

To measure mannose-sensitive agglutination of yeast, the method developed by Roe *et al*. was adapted for quantification by spectrophotometry (68). Single isolated STm colonies were used to inoculate 5 mL of LB supplemented with antibiotics and arabinose as needed and then incubated statically (or with agitation [225 rpm] for Figure S3) at 37°C for 48 hours. Cultures were centrifuged at 6,000 x g for 3 minutes to pellet cells and the supernatant was discarded. Turbidity of the cultures was measured via spectrophotometry and the OD_600_ of each strain was standardized to a value of 2.0 using LB. Yeast from *Saccharomyces cerevisiae* was obtained from Sigma-Aldrich, solubilized in Ultrapure distilled water (Invitrogen, Waltham, MA) at a concentration of 200 mg/mL using a BeadMill homogenizer (5 m/s for 2 min), and then diluted to 40 mg/mL in lukewarm distilled water. A solution of 12% (w/v) D-(+)-mannose (Sigma-Aldrich) in distilled water was prepared and filter-sterilized using a 0.2 micron syringe filter (Corning, Glendale, AZ). Finally, the STm cultures, yeast solution, and mannose solution were combined at a ratio of 2:1:1 for a final volume of 1 mL per well of a 12-well tissue culture plate or 500 µL per well of a 24-well tissue culture plate (Corning). Absorbance at 600 nm (OD_600_) was measured with a SpectraMax iD3 plate reader (Molecular Devices, San Jose, CA). The change in optical density (ΔOD_600_) for each strain was calculated by subtracting the OD_600_ values from those of the media (with and without mannose) alone, STm alone, and yeast alone.

### ELISA and dot blot

Sal4-binding assays were performed as previously described (**32**). For Sal4 ELISAs, STm strains grown as stated above and 100 μL of washed cultures (OD_600_ = 1.0) was added to a well of an Immulon 4HBX 96-well plate (ThermoFisher Scientific). The plate was covered with a plastic lid and incubated at 4°C overnight (∼18 h). The next morning, the volume in the wells was replaced with 200 μL of blocking solution (2% goat’s serum in PBS containing Tween-20 [0.1% v/v]) and the plate was incubated on a plate rocker (VWR, Radnor, PA) for 2 h at room temperature. Plates were washed three times with PBS-T, then Sal4 IgA (diluted in blocking solution) was added to each well and the plate was incubated for 1 h at room temperature on a plate rocker. After washing again three times, goat-anti-mouse IgA-HRP secondary antibody (Sigma-Aldrich, St. Louis, MO) was diluted in blocking solution at a concentration of 1:2000 and then 100 μL was added to each well. The plate was incubated for one hour at room temperature on a plate rocker, washed three times with PBS-T, and then developed with SureBlue TMB 1-Component Microwell Peroxidase Substrate (100 µL per well) (SeraCare, Milford, MA). The peroxidase reaction was stopped using an equal volume of 1M phosphoric acid and absorbance at 450 nm (Abs_450_) was measured by a SpectraMax iD3 plate reader (Molecular Devices, San Jose, CA).

For Sal4 dot blots, STm colonies from freshly streaked agar plates were used to inoculate individual wells of a 96-well microtiter plate each containing 200 μL of LB media (CELLTREAT Scientific Products, Pepperell, MA). Plates were incubated for 3 h at 37°C and 220 rpm, then 3 μL from each well was spotted onto a nitrocellulose membrane (Bio-Rad) and allowed to dry for at least 30 minutes in a fume hood at room temperature. The membrane was then submerged in blocking solution and incubated on a plate rocker overnight at 4°C. The membrane was washed 3 times in 0.1% PBS-T for 10 min prior to addition of 10 μg/mL Sal4 IgA diluted in blocking solution. The membrane was incubated for 90 min at room temperature on a plate rocker and washed again three times in PBS-T (10 minutes each). The membrane was incubated with goat-anti-mouse IgA-HRP secondary antibody (Sigma-Aldrich) diluted in blocking solution at 1:2000 for 1 h and then washed five times in PBS-T. Finally, SureBlue TMB 1-Component Microwell Peroxidase Substrate (SeraCare, Milford, MA) was applied to the membrane to detect Sal4 binding. The membrane was then imaged using a Gel Doc XR Gel Documentation System (Bio-Rad).

### Soft agar motility assay

Motility assays were performed essentially as described (25). Liquid LB media with and without Sal4 IgA was combined with an equal volume of liquified 0.6% LB agar and then poured into a 100 x 15 mm square grid petri dish (ThermoFisher Scientific). The resulting mixture was allowed to solidify at room temperature prior to inoculation with 1 μL of an STm overnight culture at the crosshair of four adjacent squares and then incubated at 37°C until bacterial growth approached the outer border of the 2x2 grid. Plates were imaged using a Gel Doc XR Gel Documentation System (Bio-Rad, Hercules, CA) and the bacterial migration diameter was measured using Fiji software version 2.9.0 (69).

### HeLa cell maintenance and invasion assay

HeLa cells were obtained from ATCC and maintained in Dulbecco’s Modified Eagle Media (DMEM) with 10% fetal bovine serum at 37°C and 5% CO_2_. The HeLa cell invasion assay was performed essentially as previously described (Deal et al., 2023). Cells were seeded at 5 x 10^5^ cells/mL in opaque 96-well tissue culture plates and grown for 24 hours. Cultures of WT (SL174), Δ*fimW::kan* (SL164), Δ*oafA::kan* (SL180), and *attTn7::kan* (SL239) were grown in 5 mL LB containing 50 µg/mL kanamycin overnight at 37°C and 225 rpm and then subcultured fifty-fold the following morning until mid-log phase was reached. Strains were pelleted at 6,000 x g for four minutes and then washed twice with 1X PBS pH 7.4. The turbidity of each strain was measured with a spectrophotometer and the OD_600_ of each strain was standardized to a value of 0.7 in PBS. Strains lacking expression of *lacZ* (SL164, SL180, and SL239) were each mixed 1:1 with SL174 (*lacZ^+^*) and then each mixture was diluted tenfold in Hank’s Balanced Salt Solution (HBSS, Wadsworth Center Media Core). The diluted mixtures were either treated with 15 ug/mL Sal4 IgA2m1 recombinant protein (Moderna, Inc.) or left untreated for 15 minutes at 37°C (28). HeLa cells were washed three times with serum-free DMEM and then the STm mixtures were applied to the monolayers. An aliquot of each mixture and treatment group was serial diluted and plated on LB agar containing 100 µg/mL kanamycin and 40 µg/mL X-gal to determine the CFU input of each strain. The culture plates were centrifuged for ten minutes at 1,000 x g (with the plate rotated 180° after five minutes to promote HeLa cell-bacteria adherence and then incubated at 37°C for one hour. Cells were washed three times with HBSS and treated with gentamicin (40 μg/mL in serum-free DMEM) to eliminate extracellular bacteria. Finally, cells were washed with HBSS and lysed with 1% Triton X-100 diluted in Ca^2+^ and Mg^2+^-free PBS. The resulting suspension was serially diluted and plated on LB agar with 100 µg/mL carbenicillin and 40 µg/mL X-gal and incubated overnight at 37°C. CFUs of each group were counted, and the competitive index [(%test strain recovered/%WT-*lacZ* recovered)/(%test strain inoculated/%WT-*lacZ* inoculated)] value was calculated for each strain and treatment group.

### Crystal violet (CV) assay

CV assays were performed as previously described with some modifications (22). Indicated strains of STm 14028s were grown to mid log phase (OD_600_ ∼ 0.7) at 37°C with aeration. Cells were then collected by centrifugation at 6,000 x g for four minutes, washed once in LB, and the centrifugation step was repeated. Turbidity of each strain was measured and then standardized to an OD_600_ of 1.0 using LB. Then, cultures were transferred to a borosilicate 16 x 125 mm glass tube and mixed 1:1 with LB medium containing Sal4 IgG at a final concentration of 15 μg/mL in a total volume of 5 mL. Tubes were incubated for 1 h at 23°C with agitation (200 rpm) and then photographed. The media was aspirated from each tube and biofilms were heat-fixed at 60°C for one hour prior to staining with 0.1% (w/v) crystal violet for 5 minutes. Tubes were rinsed once with distilled water for five minutes and the CV stain was dissolved with 30% acetic acid for 5 minutes. For quantification, 1 mL of acetic acid containing solubilized CV from each tube was divided across five wells of a clear flat-bottom 96-well plate and the absorbance at 570 nm (Abs_570_) was measured with a SpectraMax iD3 plate reader.

For IgA-induced biofilm experiments, cultures were prepared as stated above. Washed and diluted cultures were transferred to a 12-well tissue culture plate and diluted 1:1 into LB medium containing Sal4 IgA at a final concentration of 15 μg/mL in a final volume of 1 mL. Plates were incubated for 1 h at 23°C with agitation (200 rpm). The media was aspirated from each well and biofilms were heat-fixed at 60°C for one hour prior to staining with 0.1% (w/v) crystal violet for 5 minutes. Plates were rinsed once with distilled water for five minutes and the CV stain was dissolved with 30% acetic acid for 5 minutes. For quantification, the absorbance at 570 nm (Abs_570_) of the plates was measured with a SpectraMax iD3 plate reader.

### Statistical analysis and graphics

Statistical analyses were performed using GraphPad Prism 10.2.3 software (San Diego, CA). Analysis of the CRISPRi screen NGS dataset was performed using Python 3.7 (Wilmington, DE), R 4.2.1 (Vienna, Austria), and RStudio 2022.07.0 with the readxl, dplyr, tidyverse, and matrixStats packages (70–74). Figure 8 was created using BioRender.com (Toronto, Canada).

## ACKNOWLEDGEMENTS

We gratefully acknowledge the Wadsworth Center’s Applied Genomic Technologies Core for Sanger and Next-Generation Sequencing services and the Media and Tissue Culture Core for providing bacterial growth media. We would like to thank Carol Smith and Dr. Joseph Wade for their guidance regarding the experimental logistics of the CRISPRi library. We thank Dr. Cailin Deal (Moderna, Inc.) for providing recombinant Sal4 IgA2m1. SL would like to acknowledge Dylan Ehrbar for initial advice on the NGS dataset analysis and members of the Mantis lab for helpful discussions. This work was supported by the National Institutes of Health award R21AI154680. The funders had no role in study design, data collection and analysis, decision to publish, or preparation of the manuscript.

